# Fibronectin improves the impaired calcium signaling and myofilament activation in diabetic cardiomyocytes

**DOI:** 10.1101/2022.11.15.516690

**Authors:** Xin Wu, Jerome P Trzeciakowski, Gerald A Meininger, Mariappan Muthuchamy

## Abstract

Ventricular remodeling is one of the primary adaptive mechanisms in response to long-term mechanical overload in diabetes. In addition to cardiomyocyte hypertrophy, alterations in noncardiomyocyte compartments [e.g. extracellular matrix (ECM)] are an essential process in the remodeling of ventricle during diabetes. Integrins that link the ECM and intracellular cytoskeleton function as mechanotransducers to translate the mechanical force to intracellular signals. We hypothesize that mechanotransduction mechanisms are altered in diabetic cardiomyopathy mouse hearts. To test this hypothesis, force and intracellular calcium ([Ca2+]i) measurements on papillary muscle fibers were investigated in adult mouse cardiomyocytes from normal (non-db) and type 2 diabetic (db/db) mice. In addition, atomic force microscopy (AFM) was used to measure adhesion force between integrin receptors and ECM protein fibronectin (FN) by quantifying the unbinding force required to break FN-cardiomyocytes (integrin) bonds. In db/db mice, the peak active force decreased at 71% or 73% while the peak of [Ca2+]i decreased at 64% and 68% at 1 Hz or 2 Hz. In the presence of the FN (35 nM), active force was increased significantly by 40-50% in db/db mice. Furthermore, increased active force in the presence of FN was associated with 26-42% increase in [Ca^2+^]_i_ at all giving stimulations of 1 Hz and 2 Hz in db/db mice, respectively. The increased effects on force and [Ca^2+^]_i_ caused by FN were greater in ventricular muscles from db/db mice than from non-db mice. The unbinding force between FN (2.7 μM) coated AFM probes and cardiomyocyte in db/db was 52% higher than non-db (58.3 ± 0.3 pN *vs* 38.6 ± 0.9 pN. p < 0.05). The binding probability of FN-cardiomyocytes, calculated as number of force curves with adhesion / number of total force curves sampled, was significantly reduced by 30% in db/db cardiomyocytes when compared to normal. In addition, the cell stiffness, representing changes in Ca2+ signaling and cytoskeletal reorganization, was 19% increase in db/db cardiomyocytes. The presented data indicate that dynamic changes of the mechanical properties of integrin-ECM interactions may contribute to impaired intracellular Ca2+ signaling and myofilament activation in the diabetic cardiomyopathy.

## Introduction

People suffering from diabetes have imbalanced insulin production ensuing hyperglycemia that if left untreated can cause cardiovascular disease, kidney failure, nerve damage, blindness, and ultimately death. About 90% of all cases of diabetes are non-insulin-dependent diabetes mellitus (DM), and are called type 2 DM. The cardiac complications associated with Type 2 DM are due to both increased coronary heart disease secondary to atherosclerosis and a specific diabetic cardiomyopathy resulting in ventricular dysfunction (23, 42, 66, 78). In addition to cardiac ventricular myocytes (CVM) hypertrophy during mechanical overload, alterations in non-CVM compartments (e.g. extracellular matrix (ECM)) are an essential process in the remodeling of ventricle in diabetes (6, 36). During progress of diabetes, there is an increase in expression and deposition of fibronectin (FN) and collagen (CN) in non-CVM compartments (4, 27, 28, 47, 49, 60, 67).

Integrins are a large family of transmembrane glycoproteins that provide a connection between the intracellular cytoskeleton and ECM. Of the approximately 24 known integrins, CVMs express at least 4 prevalent subtypes, which include: α1β1, α3β1, α5β1 and α7β1. α3β1 can bind to CN, FN and laminin, whereas, α5β1 binds to FN; α1β1 and α7β1 bind to laminin (58). α1β1, α3β1 and α5β1 recognize ECM proteins containing RGD sequence motifs, which are present in FN (62, 64, 100). Furthermore expression of β3 and β5 integrins has also been reported in CVM (50, 55, 74). FN is a large glycoprotein found in plasma, in extracellular matrix, and on cell surfaces. It promotes cell-cell and cell-matrix interactions and thus plays an important role in tissue construction and reconstruction. A key observation from our previous studies is that at least 3 different integrins regulate voltage-gated L-type Ca2+ channels (CaL), Ca2+ activated K+ channels and myogenic responses in vascular systems, cardiomyocytes, neuronal cells, and in heterologously expressed neuronal, smooth muscle and cardiac CaL in HEK cells (35, 90, 94, 95, 97, 100, 102, 108). In mouse ventricular fibers, RGD peptide and collagen modulated the force production partly associated with changes of intracellular Ca^2+^ concentration ([Ca^2+^]_i_) secondary to activation of α5β1 integrin (64). However, fibronectin increased force production associated with increased activity of L-type calcium channel, [Ca2+]i, and PKA phosphorylation (94). Using AFM in our laboratory, we have reported that the changes of mechanical properties by FN-CVM, FN-vascular smooth muscle cells and FN-endothelial cells interactions were primarily by activation of α5β1 integrin (76, 81, 100).

Numerous studies have been indicated that alterations in CaL and [Ca2+]i are a primary stimulus for the cardiac hypertrophic response in diabetes. Diabetic cardiomyopathy is characterized by myocyte damage and myocardial fibrosis, leading to decreased elasticity and impaired contractile function. Several studies have suggested that the existence of a diabetic cardiomyopathy is independent of preexisting coronary vascular sclerosis and arterial hypertension. The incidence of congestive heart failure in diabetic patients is 2.5-5 fold increased and is independently of age, arterial hypertension, and coronary artery disease (45). Imbalance in the production and the degradation of ECM proteins may lead to structural alterations including basement membrane thickening and ECM protein deposition in the tissues of diabetes (67, 70). FN mRNA and proteins are increased in hearts from diabetes or hyperglucose treated rats and mouse, and the increase of FN expression might be related to leptin and endothelin 1 (53, 60, 70, 71). Circulating cellular FN was also significantly elevated (3 – fold) in patients with diabetes (46). ECM proteins (e.g. CN and FN) deposits compromise tissue compliance and causing myocardial stiffness (8, 14). Kinnunen group found a tight connection between lipids and amylin aggregates deposited on lipid bilayers using fluorescence resonance energy transfer microscopy (21). This procedure can cause membrane structural changes. There are some studies shown decreased Ca^2+^ activities and cardiac contractilities from in vivo and isolated working heart in diabetes (38, 69, 105).

However, there is no studies on type 2 DM using atomic force microscopy (AFM) and a few studies on papillary muscle force-[Ca^2+^]_i_ relationship. It has been well demonstrated that cardiac contractile and [Ca^2+^]_i_ dysfunction in insulin-deficient (type 1) diabetic animals with isolated working hearts and papillary muscles (17, 24, 40, 56, 69, 86). Relatively few studies of cardiac contractility and Ca^2+^ signaling have been conducted with type 2 DM (8, 9, 11).

In this study we examined the relationship between extracellular matrix protein and cardiac muscle contractility, and mechanotransduction characteristics such as adhesion force and cell stiffness in type 2 diabetic mouse heart.

## Methods and Materials

### Adult CVM preparation

Mouse (12-20 weeks, type 2 diabetes mellitus (homozy db/642, db/db, C57BLK6 strain) myocytes were prepared as described (100). The mice were held in an environmentally controlled institutional animal facility under a 12 hour/12 hour light/dark cycle. All procedures were performed in strict compliance with the guidelines of National Institutes of Health Guide for the Care and Use of Laboratory Animals under a protocol approved by the university’s Institutional Animal Care and Use Committee.

The hearts from adult male mouse were harvested under anesthetic conditions and put into ice-cold Ca^2+^ free physiological saline solution (PSS) containing (in mM): 133.5 NaCl, 4 KCl, 1.2 NaH2PO4, 1.2 MgSO4, 10 HEPES, and 11 glucose, pH 7.4. 10 mM 2,3-butanedione monoxime (BDM) was present during dissecting procedure. The aorta was cannulated and the heart was mounted in a Langendorff perfusion system with PSS containing 25 μM Ca2+ and selected collagenases. Then heart was removed and transferred to a Petri dish at PSS with 100 μM Ca2+. The ventricle muscles were cut into small pieces. The pieces were gently triturated using a fire-polished Pasteur pipette to release single cells. The collected cells were then re-suspended in the PSS containing 200 μM Ca^2+^. After this procedure, the cells were stored at room temperature and used within 6 hours. All experiments were carried out at room temperature (22–23°C).

A suspension of freshly dispersed cells was plated onto a thin dish on the stage of an inverted microscope for at least 30 min in PSS with 1.8 mM Ca2+ before AFM experiment.

#### Application of atomic force microscopy (AFM)

The force contact mode operation was used for measurements of adhesion force (unbinding force) on retraction curve and relative cortical membrane stiffness (elasticity) on approach curve using a Bioscope AFM system (Model 3A, Digital Instruments, Santa Barbara, CA), which was mounted on an Axiovert 100 TV inverted microscope (Carl Zeiss, Germany), in ventricular cells (98, 100, 101). The AFM probes used were silicon nitride microlevers with conical tips (model MLCT-AUHW, Santa Barbara, CA). Tip radius were < 20 nm and mean spring constant was approximately 14.4 ± 0.6 pN/nm. AFM probes were labeled with FN (1 mg/ml). Polyethylene glycol (PEG) was used to cross-link proteins onto AFM probes at room temperature. For each experiment, the position of the protein (e.g. FN) labeled probe was controlled to repeatedly touch and retract (*Z*-axis) from the cell surface. Force curves were recorded for these repeated cycles of probe approach and retraction at 0.5 Hz scan frequency and a Z-axis movement of 800 nm.

#### Force–[Ca^2+^]_i_ measurements in intact papillary muscles

Right ventricular papillary muscle bundles were extracted and mounted as previously described (79, 94). The muscle measurement equipment suite provided all the optics and electronics needed for measuring [Ca^2+^]i using Fura-2 AM dye. Measurements were collected through a different data acquisition suite (National Instruments A/D board and LabVIEW software) with the digital oscilloscope suite providing continuous monitoring. A mercury lamp and filter wheel provided alternating ultraviolet (UV) pulses of 340 nm and 380 nm at 250 Hz with pulse duration of 1.5 ms to illuminate the bundle. The combinations of microscope, dichroic mirror, filter and photomultiplier tube collected the Fura-2 fluorescence. A synchronized electronic integrator parsed and averaged the fluorescence from both 340 nm and 380 nm illuminations to the A/D system. The loading solution consisted of Krebs–Henseleit (KH) with 10 μM Fura-2 AM, 4.3 mg l-1 N,N,N’,N’-tetrakis (2-pyridylmethyl) ethylenediamine, and 5.0 g/L cremophor. The KH to dimethyl sulfoxide volume ratio of the loading solution was 333:1. A loading duration of 1.5 h with 20 min of de-esterification gave signals of greater than 3-fold over background fluorescence. The ratio, R, of fluorescence from 340 nm excitation to fluorescence from 380 nm excitation was calculated after subtracting background fluorescence. Calcium concentration was calculated using the equation with Kd equating to Kd x β after subtracting background fluorescence (34):

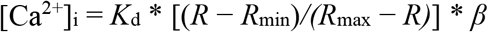

β is the ratio of the 380 nm signal at zero calcium versus the 380 nm signal at saturating calcium (39.8 μM). The ratio of 340/380 fluorescence was converted to [Ca^2+^]_i_ using a standard method (34). Minimum ratio (R_min_) was determined in zero Ca^2+^ with 10 mM EGTA, and maximum ratio (R_max_) was determined in 39.8 μM Ca^2+^. The experimental protocols were the same as described in the earlier section.

From the calcium and force values, the maximal force, time-to-peak Ca^2+^ amplitude, time-to-peak force, the time from peak of [Ca^2+^]_i_ to peak of force, and the maximum rate of isometric tension development [+d*F*/d*t,* (mN/mm^2^)/s)] present as properties of contraction. 50% decay time from the peak of Ca^2+^ and 50% relaxation time from the peak of force, and the maximum rate of relaxation [-d*F*/d*t,* (mN/mm^2^)/s] represented as relaxation properties. The +d*F*/d*t* and −d*F*/d*t* was calculated using the following equation, where *F(t)* is the measured force at a particular time *t* and *ΔT* is the 1 ms sampling period:

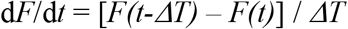

Averaging a three data point window for each single d*F*/d*t* calculation minimized the noise. Data analyses were done as described as our previous publications (30, 80, 94). We analyzed the force–calcium values at specific points during a twitch cycle. The points consisted of: (1) the resting point ‘*a*’, (2) maximum calcium point, ‘*b*’, and (3) maximum force point, ‘*c*’. Active force, [Ca^2+^]_i_ and active force/delta gain of [Ca^2+^]_i_ were calculated at points *a*, *b* and *c*.

#### General data analysis

All data are expressed as means ± SEM. Statistical analyses were done using either a Student’s paired t test or a two-way ANOVA with Fisher’s or Bonferroni/Dunn’s post hoc test. Repeated measures ANOVA were used for comparison of repeated measurements within the same group (e.g. response to increasing pacing frequency). P <0.05 was regarded as statistically significant.

## Results

### General characteristics of non-db and db/db mice

At 12–20 week of age, db/db mice weighed significantly more than their lean nondiabetic (non-db, C57BLK6 strain) littermates (Table 1). This increase in body weight was primarily due to increased fat deposition in abdomen and internal organs that can be observed during experiments. The differences in heart weight, waist size, liver weight and kidney weight were significant between the two groups. The ratios of heart weight and body weight were significant lower than non-db group. In addition, db/db mice had significantly elevated fasting serum levels of glucose, indicating their diabetic status with hyperglycemia (Table 1).

**Table 1.**
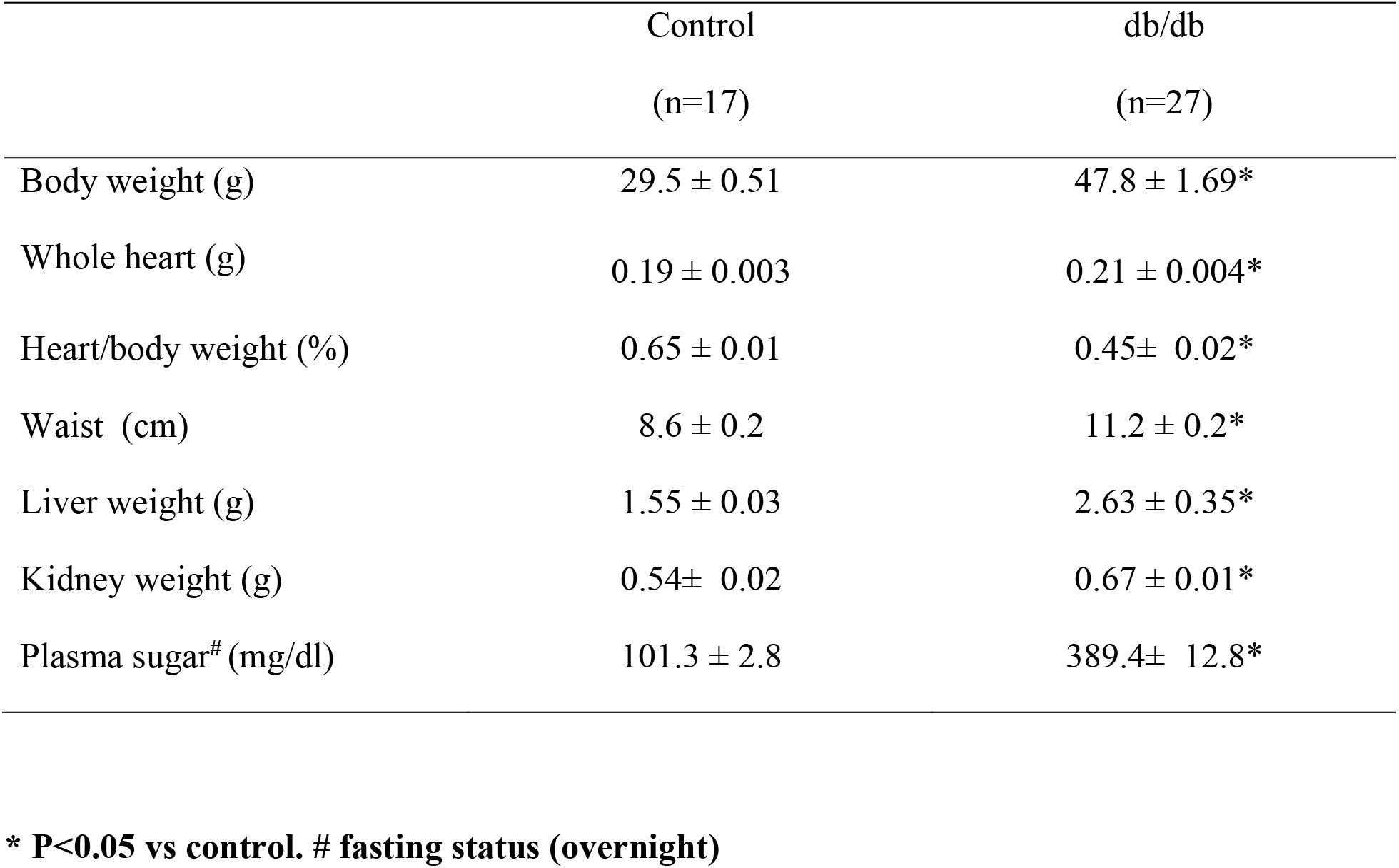
General characteristics of control and diabetic (db/db) mice.

|  | Control | db/db |
| --- | --- | --- |
|  | (n=17) | (n=27) |
| Body weight (g) | 29.5 ± 0.51 | 47.8 ± 1.69* |
| Whole heart (g) | 0.19 ± 0.003 | 0.21 ± 0.004* |
| Heart/body weight (%) | 0.65 ± 0.01 | 0.45± 0.02* |
| Waist (cm) | 8.6 ± 0.2 | 11.2 ± 0.2* |
| Liver weight (g) | 1.55 ± 0.03 | 2.63 ± 0.35* |
| Kidney weight (g) | 0.54± 0.02 | 0.67 ± 0.01* |
| Plasma sugar<sup>#</sup> (mg/dl) | 101.3 ± 2.8 | 389.4± 12.8* |

**\* P<0.05 vs control. # fasting status (overnight)**

### The db/db hearts exhibit decreased contractile function

To explicate how diabetes affects heart function in tissue level, we measured the force and [Ca^2+^]_i_ simultaneously in the right ventricular papillary muscle fibers at room temperature. The Ca^2+^ transients and force measured at 1 Hz in papillary muscles from non-db mice (n=7) and db/db mice (n=10) are shown in figures 1A. Active force that describes the difference between the maximum and minimum force (passive tension), developed by the ventricle fibers decreased at 1 Hz stimulating rate in db/db mice (Figure 1). The time to peak of [Ca^2+^]_i_ (Figure 1A) and time to peak force (Figure 1B) were not significantly changed in the ventricle muscle from db/db mice. The relaxation time (50% from the peak of the force) were 29% and 26% slower in papillary muscles from db/db mice (76 ± 7 ms at control vs 98 ± 6 ms at db/db at 1 Hz; 63 ± 9 ms at control vs 79 ± 5 ms at db/db at 2 Hz. (Figure 1B), respectively. The Ca^2+^ transients decay (declined to 50 % of the peak) were 32% and 21% slower at 1 Hz (128 ± 13 ms at control *vs* 169 ± 8 ms at db/db, p < 0.05) and 2 Hz (103 ± 10 ms at control vs 125 ± 5 ms at db/db, p<0.05) in db/db mice, respectively. This latter observation suggests a possible defect in SR calcium uptake through SERCA2a (11).

**Figure 1.**
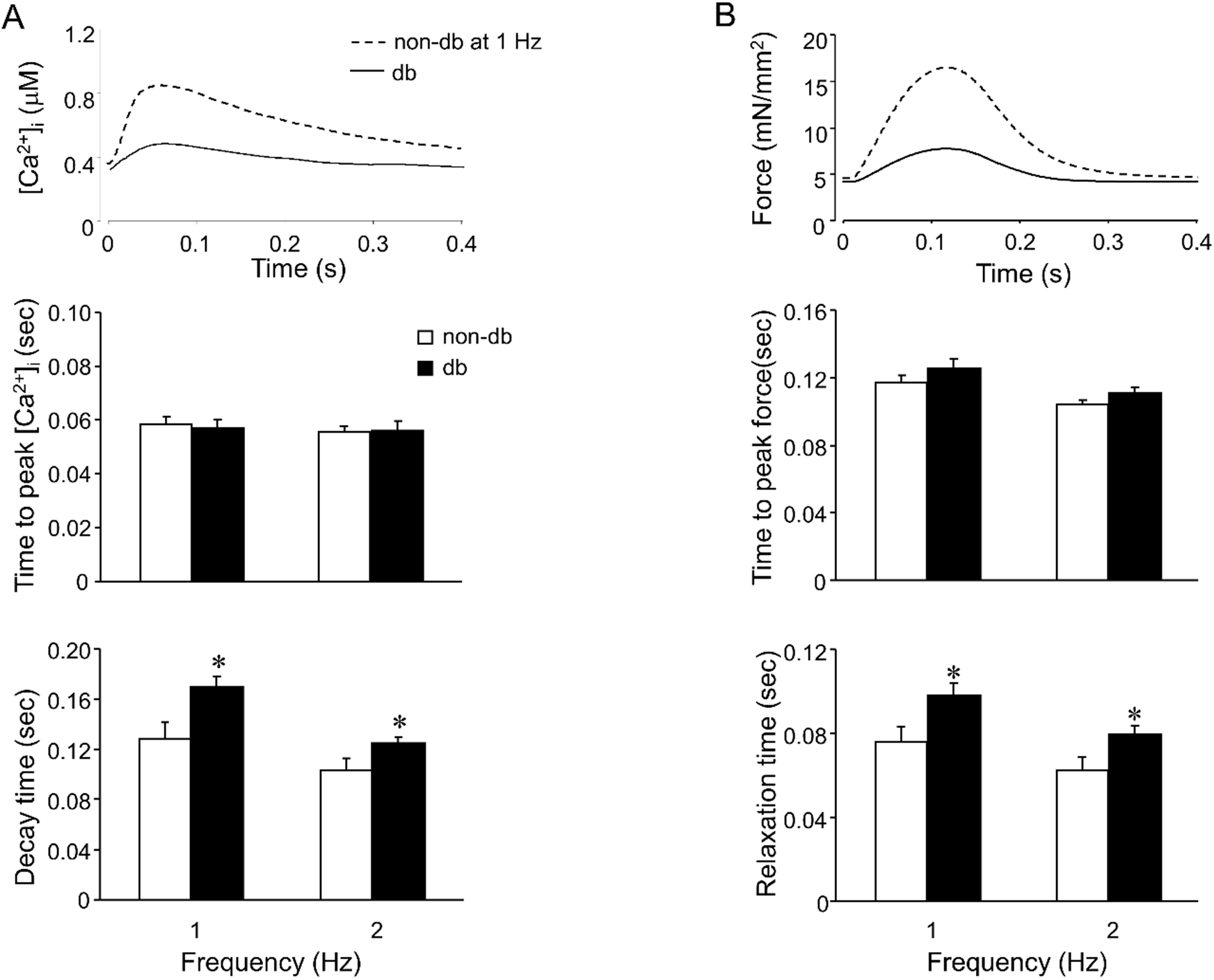
Ca^2+^ transient and force during a contraction cycle in a papillary muscle from non-db and db/db mice. Raw data showing [Ca^2+^]_i_ transients (A) and force (B) in top panels. Force and [Ca^2+^]_i_ with respect to time at a stimulation rate = 1 Hz. Decay time of [Ca^2+^]_i_ and force relaxation time are the times representing EC50 values. There are significant difference in non-db group (n=7) vs db/db mice group (n=10). Data are presented as mean ± SEM. * *p* < 0.05 *vs* non-db group.

Rates of force generation and relaxation (+d*F*/d*t* and −d*F*/d*t*) were calculated as described in Methods. Ventricle muscles from db/db mice showed significant decreased in both +d*F*/d*t* and −d*F*/d*t* at all the tested stimulation frequencies, 1 and 2 Hz (Figures 2). The maximum rate of force generation (+d*F*/d*t*) decreased from 70% at 1 Hz to 72% at 2 Hz. The rate of force relaxation decreased from 80% at 1 Hz to 81% at 2 Hz, respectively, in the ventricular muscles from db/db mice.

**Figure 2.**
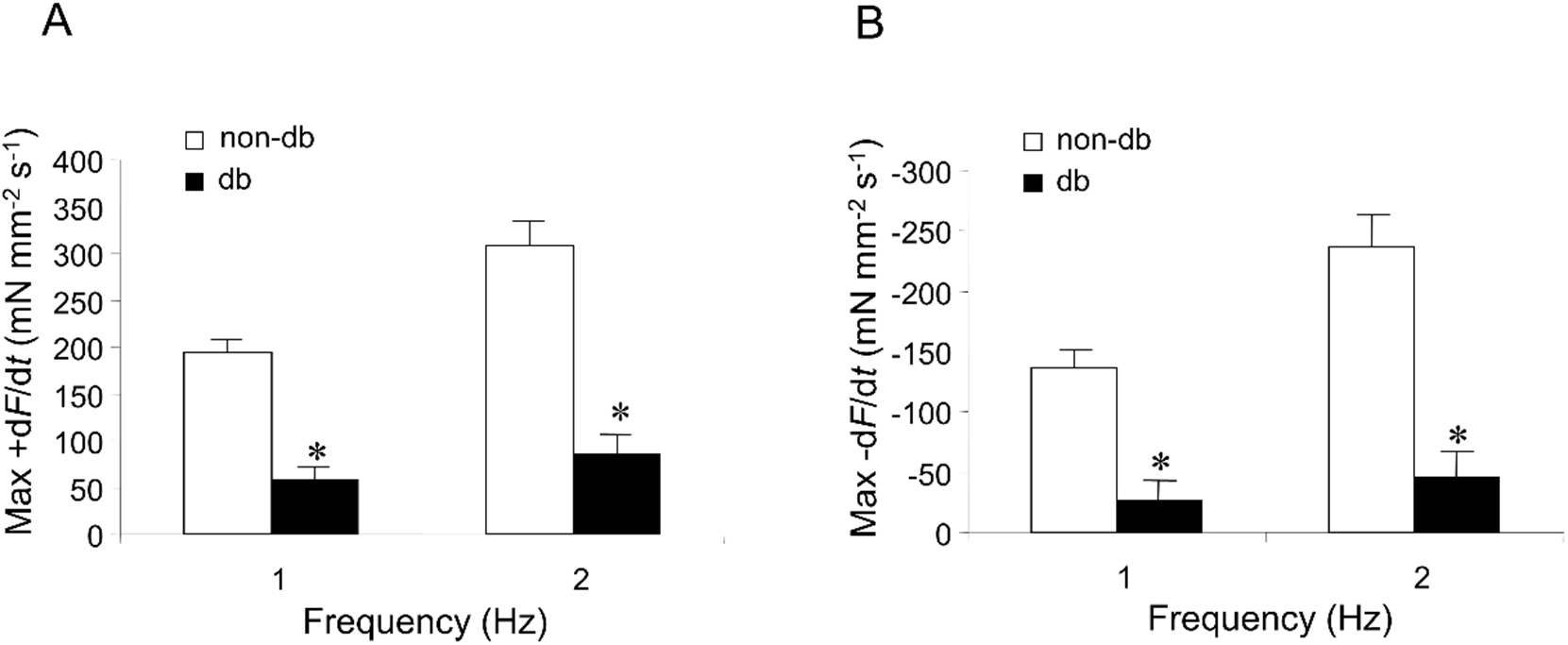
FN enhanced both the contraction and relaxation parameters in papillary muscle. (A) and (B) The maximum rates of contraction and relaxation (+d*F*/d*t* and −d*F*/d*t*) are decreased in fibers from db/db mice at stimulation of 1 Hz and 2 Hz. Data are presented as mean ± SEM. n = 7 for non-db group; n=10 for db/db group. * *p* < 0.05 *vs* non-db group.

Figure 3A shows typical force - [Ca^2+^]_i_ loops for papillary fibers before and after FN application at 1 Hz. Note three distinct points labeled as *‘a’, ‘b’* and *‘c’* on the force-[Ca^2+^]_i_ hysteresis loop. Point *‘a’* represents the resting (basal) point, point *‘b’* represents maximal [Ca^2+^]_i_ concentration and point *‘c’* represents maximal force. At point ‘*a*’, increase in stimulation frequency did not significantly change the [Ca^2+^]_i_ and diastolic force of the fibers in either the non-db or db/db mice. Figures 3B and 3C show the changes in the maximum active force, maximum [Ca^2+^]_i_, and delta gain (DG) of active force divided by the change in [Ca^2+^]_i_ (active force/Δ[Ca^2+^]_i_) for stimulation frequencies 1.0 Hz and 2.0 Hz that occur at point ‘b’ and point ‘c’, respectively. The DG of the active force/Δ[Ca^2+^]_i_ is defined as the active force divided by the difference in [Ca^2+^]_i_ from point ‘b’ to point ‘a’ or from point ‘c’ to point ‘a’. Since delta gain quantifies changes in force per unit calcium, the alteration in this parameter could represent changes in the myofilament activation processes (30, 64, 80, 94), such as an increase in Ca2+ sensitivity. The results demonstrate that at point ‘b’ (Figure 3B), the active force significantly decreased by 77% and 77% while the maximal Ca2+ was decreased by 64% and 57% at 1 Hz and 2 Hz in the ventricular muscles from db/db mice. Furthermore, DG significantly decreased by 57% and 67% at 1 Hz and 2 Hz, respectively. At point ‘c’ (Figure 3C) the maximum active force was decreased by 71% and 73% at 1 Hz and 2 Hz, respectively, in the ventricular muscles from db/db mice. The [Ca2+]i was also decreased by 61% at 1 Hz and by 56% at 2 Hz. The DG at point ‘c’ significantly decreased by 50% and 61% at 1.0 and 2.0 Hz, respectively, in the ventricular muscles from db/db mice. These results indicated db/db hearts exhibited decreased contractile function probably through decrease of [Ca2+]i.

**Figure 3.**
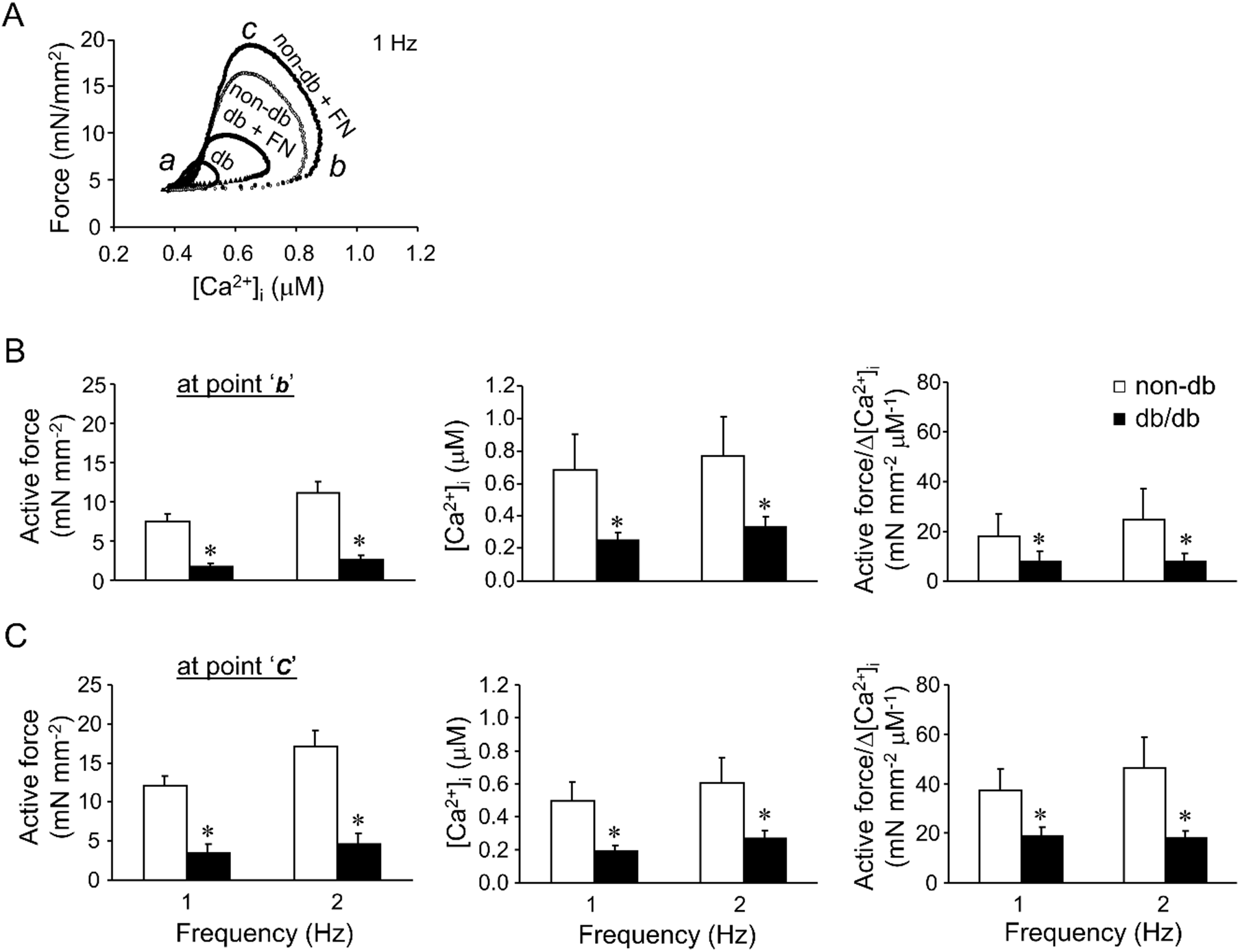
Analysis of the force-[Ca^2+^]_i_ loop in papillary muscles from non-db and db/db mice. (A) Representative force-Ca^2+^ loop in the absence and presence of fibronectin (FN) in non-db and db/db groups at a stimulation rate = 1 Hz. *‘a’* is the resting point; ‘*b’* is the peak [Ca^2+^]_i_ point; and ‘*c*’ is the peak force point. (B) and (C) [Ca^2+^]_i_, active force, and active force/Δ[Ca^2+^]_i_ in papillary muscles from non-db (n=7) and db/db mice (n=10). Data are presented as mean ± SEM. * *p* < 0.05 *vs* non-db of corresponding rates.

### Fibronectin enhanced active force and [Ca2+]i in type 2 DM

Several studies reported that diabetes induced cardiac remodeling related to ECM expression. The ECM proteins (e.g. FN) could impair the cardiac functions in DM (54, 71, 84). To determine how integrin ligands modulate the contractility in db/db mice, force-[Ca^2+^]_i_ relationship was analyzed in the presence of FN (35 nM). We have reported that FN can increase peak force and peak [Ca2+]_i_ in FVB/N strain mice (94). Here, FN also increased the peak of active force (at point ‘c’) by 34% at 1 Hz and 11% at 2 Hz in the presence of FN in ventricular muscles from *non-db* mice (Figure 4A. n=7), respectively. Peak of [Ca^2+^]_i_ (at point ‘b’) was increased at 41% at 1 Hz and 26% at 2 Hz in the presence of FN in ventricular muscles from *non-db* mice, respectively (p<0.05, Figure 4B. n=7). Meanwhile, peak of active force (at point ‘c’) was increased at 50% at 1 Hz and 43% at 2 Hz in the presence of FN in ventricular muscle from *db/db* mice, respectively. Peak of [Ca^2+^]_i_ (at point ‘b’) was increased at 89% at 1 Hz and 60% at 2 Hz in the presence of FN in ventricular muscle from *db/db* mice, respectively (p<0.05, n=10).

**Figure 4.**
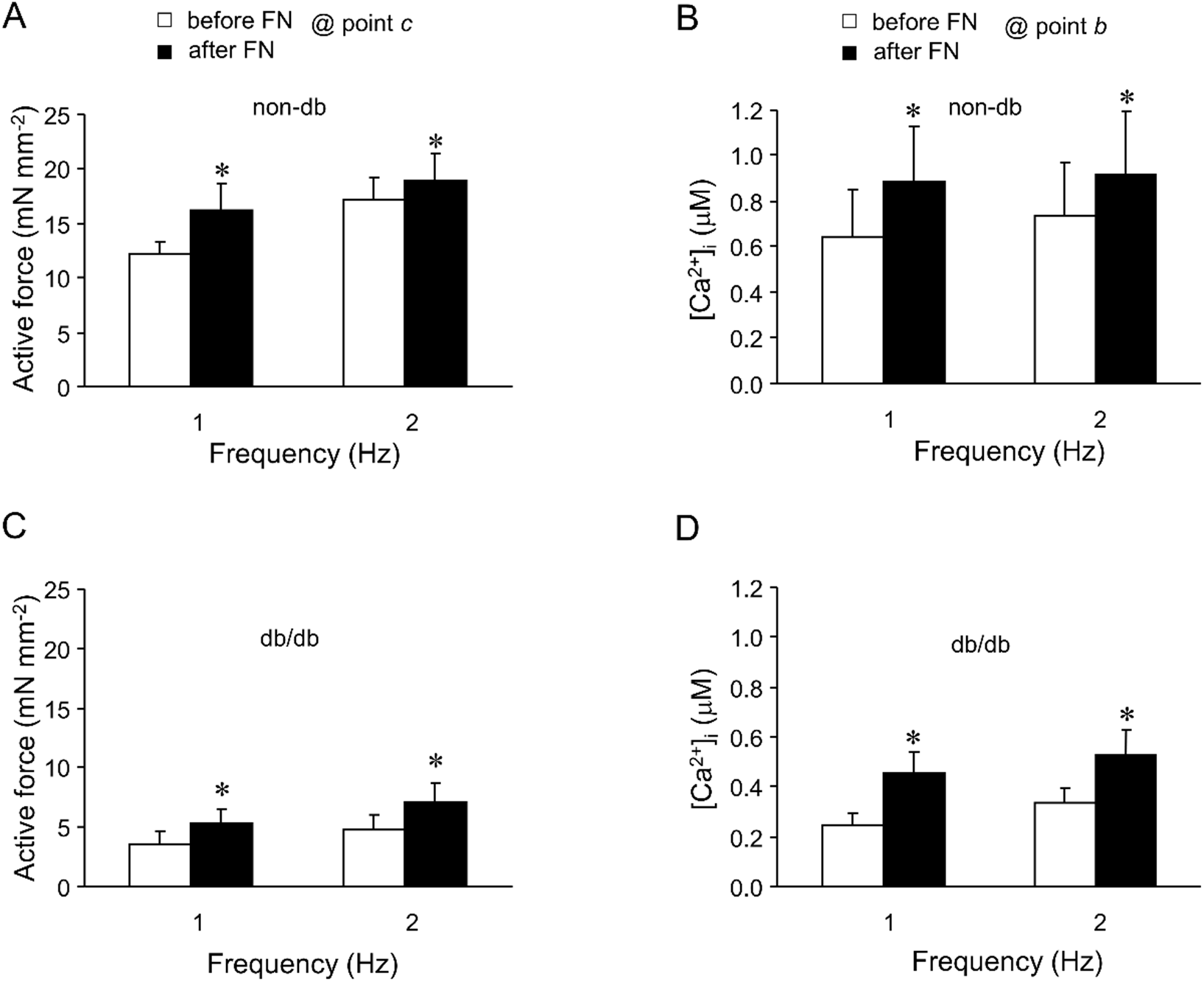
FN enhanced active force and [Ca2+]i in non-db and db/db mice. Active force (A and C, at point ‘c’) and peak of [Ca2+]i (B and D, at point ‘b’) were increased in the presence of FN in ventricular muscles from non-db (n=7) and db/db mice (n=10) at 1 Hz and 2 Hz. * *p* < 0.05 *vs* before FN application.

Ventricle muscles showed significant increased after FN application in both +d*F*/d*t* and – d*F*/d*t* at all the tested stimulation frequencies, 1 and 2 Hz in ventricular muscle from *db/db* mice (Figures 5A and 5C). The rate of force generation increased from 35% at 1 Hz to 34% at 2 Hz (Figure 5A). The time to peak of [Ca^2+^]_i_ and time to peak force were also significantly decreased in the presence of FN. The time from peak of [Ca^2+^]_i_ to peak of force was significant shorter at 1 Hz after FN (68 ± 7 ms in non-db *vs* 60 ± 6 ms in db/db, p < 0.05, n=10). The rate of force relaxation increased from 75% at 1 Hz to 76% at 2 Hz (Figure 5C). The relaxation time (50% from the peak of the force) were 19% and 18% faster (98 ± 5 ms before FN vs 80 ± 6 ms after FN at 1 Hz; 79 ± 5 ms before FN vs 65 ± 4 ms after FN at 2 Hz, p<0.05) in papillary muscles after application of FN, respectively. The Ca^2+^ transients decay (declined to 50 % of the peak) were 27% and 16% faster at 1 Hz and 2 Hz (169 ± 8 ms before FN vs 80 ± 6 ms after FN at 1 Hz; 122 ± 7 ms before FN vs 105 ± 6 ms after FN at 2 Hz, p<0.05) in the presence of FN, respectively. These results indicated fibronectin enhanced active force probably through increase of [Ca2+]_i_ in type 2 DM.

**Figure 5.**
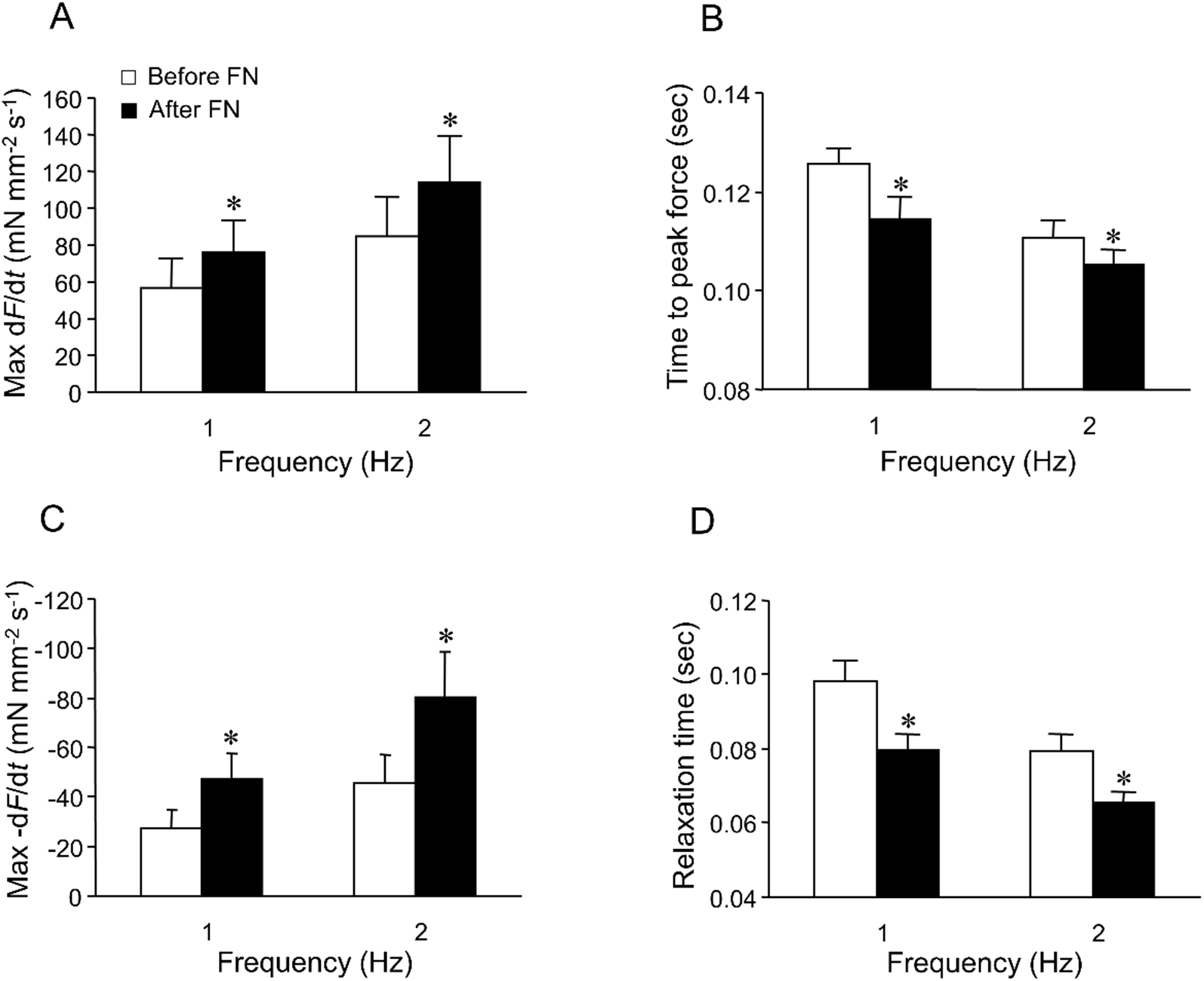
FN enhanced both the contraction and relaxation parameters in papillary muscles from db/db mice. (A) and (C) The maximum rates of contraction and relaxation (+dF/dt and −dF/dt) are enhanced in fibers treated with FN. (B and D) Time to peak force and the time to 50% off the peak of maximum force are faster in the FN-treated fibers. Data are presented as mean ± SEM. * p < 0.05 vs before FN application.

### FN enhanced ventricular active force in type 2 DM greater than in non-DM

To investigate whether there is a difference by FN on ventricular muscles from non-db and db mice, we compared the results in the presence and absence of FN from these 2 groups. Figure 6 shown a summary of FN enhanced maximum active force (normalized to the value before FN) in ventricular muscles from non-db or db/db mice. There were 4.1-fold (at 2 Hz) greater on force enhancement (comparing enhanced portion after FN application) and 2.2-fold (at 1 Hz) on [Ca^2+^]_i_ enhancement after FN application in ventricular muscles from *db/db (n=10)* than in ventricular muscles from non-*db* (n=7, p<0.05). FN also has great effects on force generation in db/db mice than in non-db mice. There were 3.1-fold greater in rate of force generation (+d*F*/d*t*) and 3.5-fold greater in rate of relaxation *(-dF/dt)* after FN application in db/db than in non-db at 2 Hz, respectively. These results indicated FN enhanced myocardium active force in type 2 DM greater than in non-DM.

**Figure 6.**
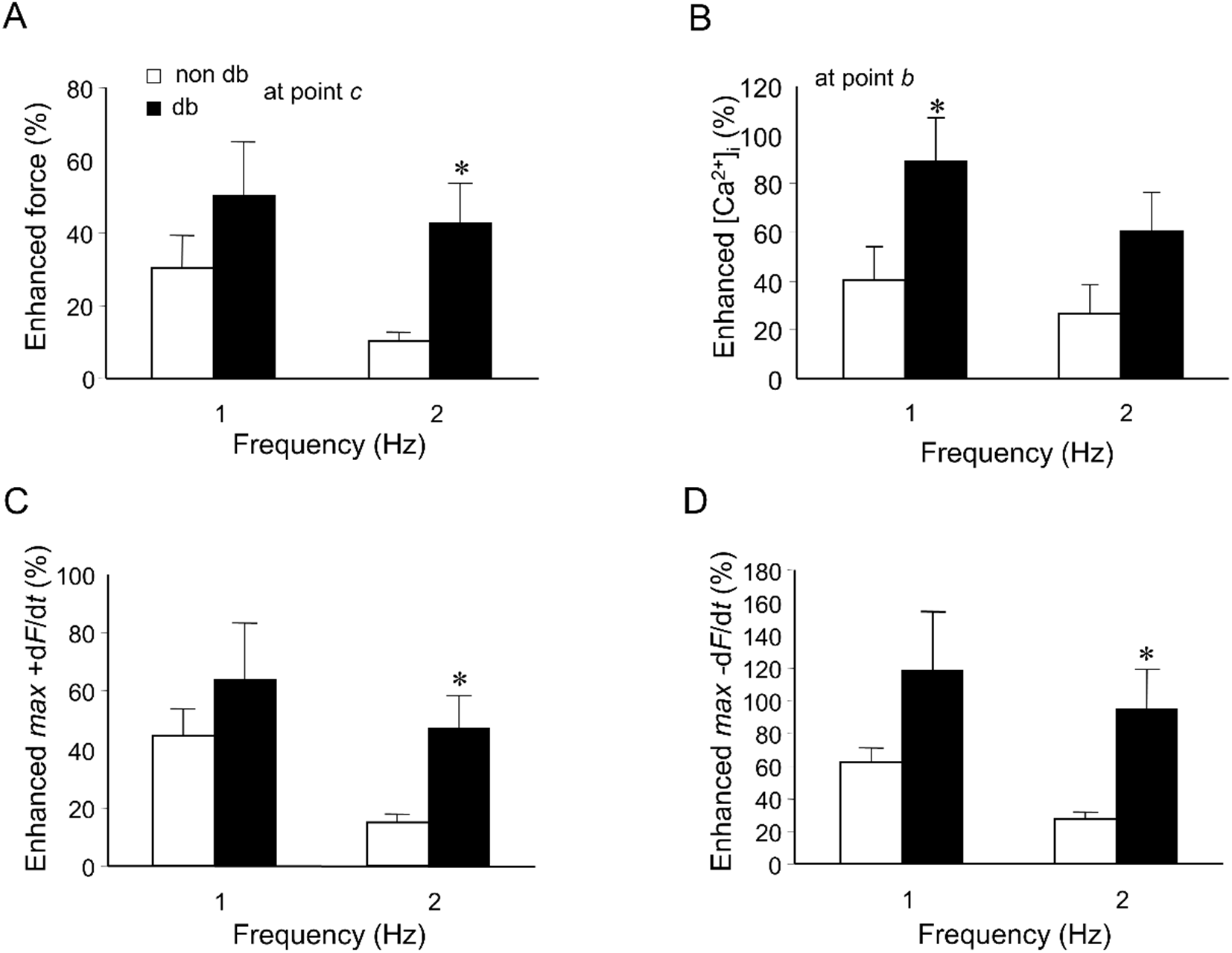
FN-enhanced ventricular contractility in db/db mice is greater than in non-db. (A) and (B) Summary of FN enhanced maximum active force (% of before FN application) and [Ca2+]_i_ in ventricular muscles from non-db and db/db mice (db). There were 4.1-fold (at 2 Hz) greater on force enhancement (comparing enhanced portion after FN application) and 2.2-fold (at 1 Hz) on [Ca^2+^]_i_ enhancement after FN application in ventricular muscles from *db/db* (n=10) than in ventricular muscles from non-*db* (n=7). (C) and (D): Summary of FN enhanced rate of force generation (% of before FN application, max +dF/dt, and rate of maximum relaxation (max −dF/dt, C) in ventricular muscles from non-db and db/db mice. There were 3.1-fold greater in rate of force generation (+d*F*/d*t*) and 3.5-fold greater in rate of relaxation (-d*F*/d*t)* after FN application in db/db than in non-db at 2 Hz, respectively. * p < 0.05 vs non-db mice.

### FN enhanced adhesion force and cell membrane elasticity in CVM from type 2 DM

To measure the cell membrane stiffness and the adhesion force (unbinding force) between FN and cardiomyocyte surface, AFM probes coated with FN were applied to the membrane surface of non-contracting cardiomyocytes as described previously (100). To analyze the distribution of adhesion events, the observed adhesion events plotted as histograms were fitted with Gaussian distributions to resolve integrin-FN bond adhesion force. The peak bond rupture force (initial peak of the adhesion force in Figure 7 and Figure 8) of FN-integrin was approximately 38.6 ± 0.9 pN in non-db mice (n=10) and 58.3 ± 0.3 pN in db/db mice, respectively (n = 6, p< 0.05), which likely represents a FN-integrin single bond unbinding force (75, 76, 81, 100). The bar at the right margin of the Figure 7A shows the probability of FN-integrin adhesion events defined as the percentage of the curves with adhesion divided by total recorded curves. Under conditions defined for these experiments, the probability of adhesion events between the FN-coated probe and cardiomyocyte was 30% reduced (62% in non-db vs 44% in db/db, Figure 7). The integrated force value determined as an average force from all adhesion events was also calculated as described in the methods section and mentioned in previous report (100). The integrated adhesion force between FN and the myocyte was 77.8 ± 1.7 pN in non-db mice. The integrated force was increased by 56% in db/db mice (Figure 8B. p<0.05).

**Figure 7.**
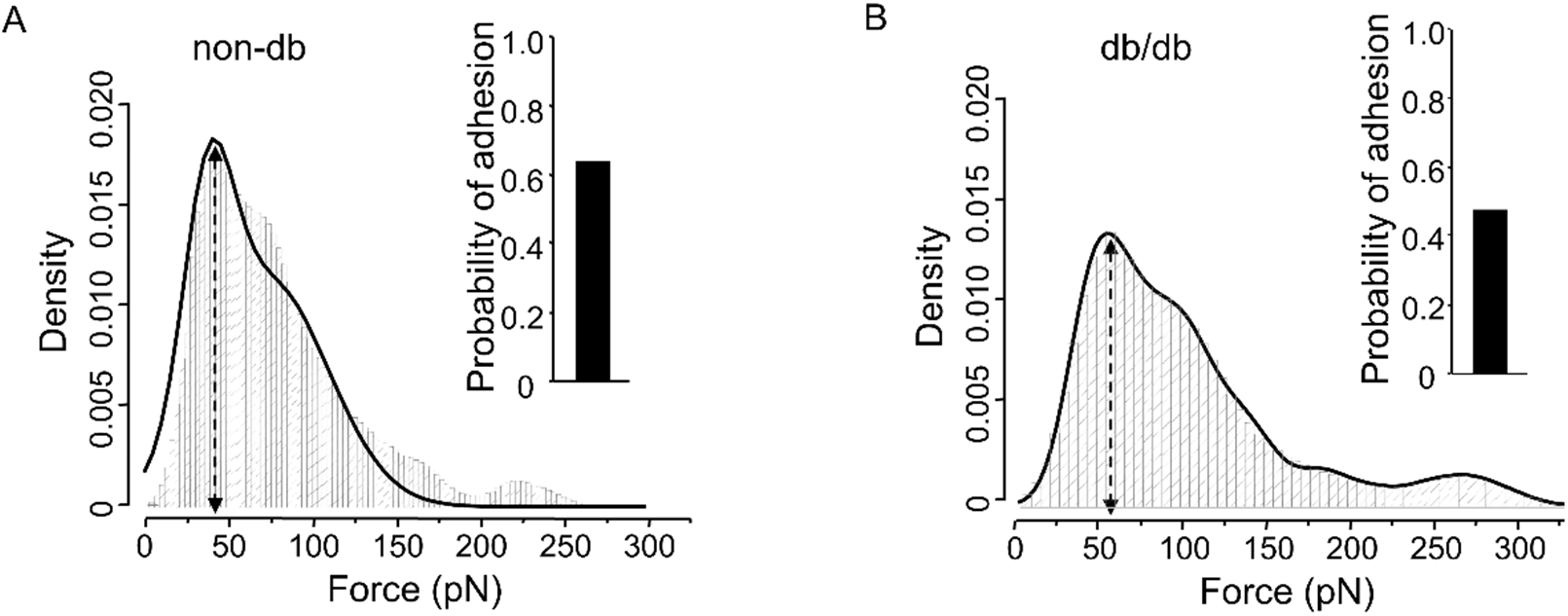
Summary of adhesion force results with FN-coated AFM probe in cardiomyocytes from non-db and db/db mice. (A) and (B): Analyses of force-density of adhesion event during FN-coated probes retraction in cardiomyocytes. Observed adhesion force and corresponding number of events in the experiments (50 curves/cell for total 300 - 500 curves) were plotted as histograms in non-db (A, n=10) and db/db mice (B, n=6). Solid lines represent the results that fitted with multiple Gaussian distributions. Insets: integrin-FN binding probabilities (solid bars) in non-db (A) and db/db (B).

**Figure 8.**
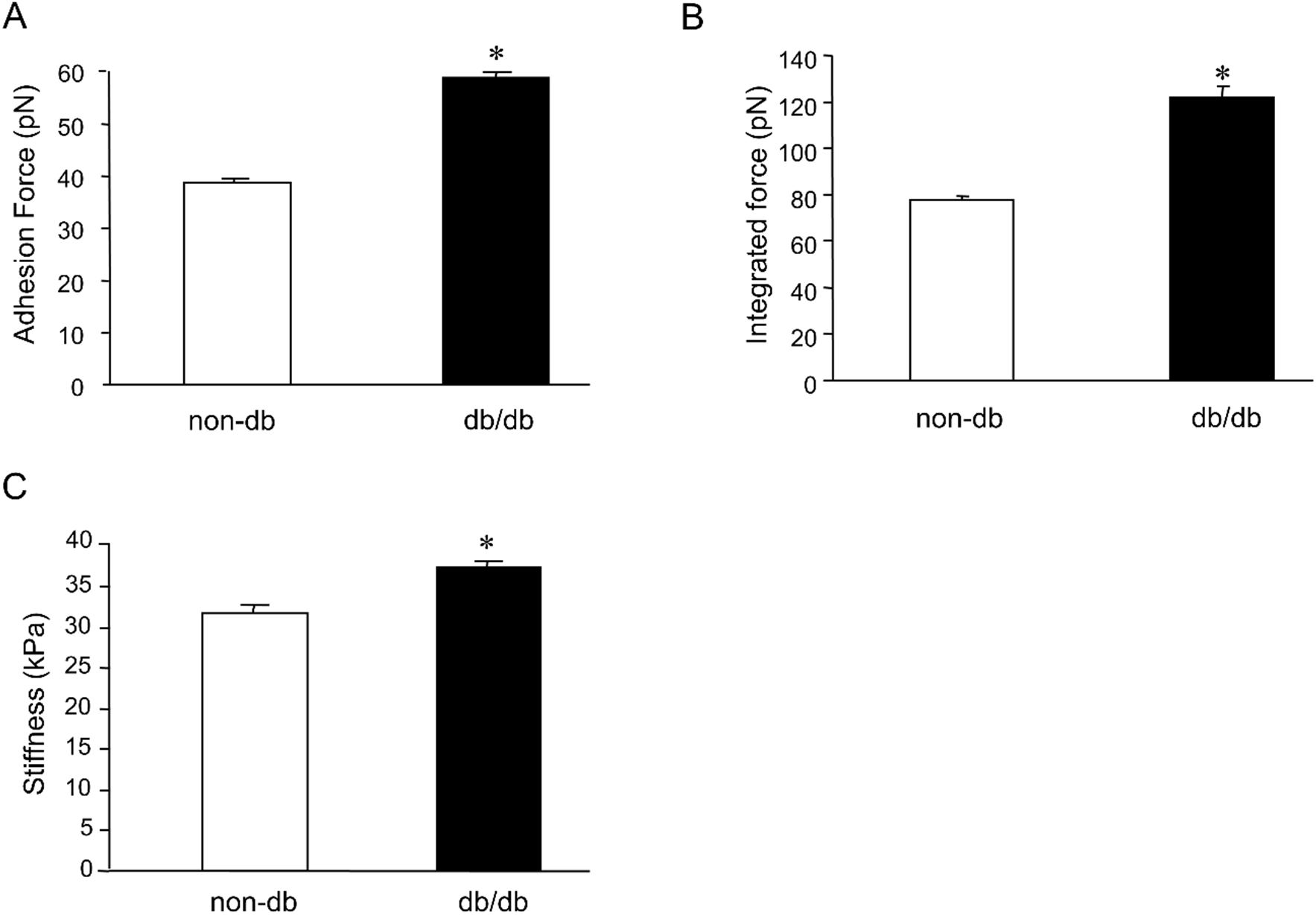
Summary results of adhesion characteristics of cardiomyocytes with FN-coated AFM tips in non-db (n=10) and db/db mice (n=6). Adhesion force (A), integrated force (B) and cell stiffness (C) were significantly increased in db/db mice. \**p* < 0.05 vs non-db group.

The cell stiffness represents changes in intracellular signaling and cytoskeletal reorganization. CSK in the depth of cytoplasm, cell membrane components (e.g. cholesterol) and its coupling to the CSK, and intracellular components other than CSK (e.g. Ca^2+^ and osmolarity) contributed to the stiffness(15, 29, 39, 41, 72, 85, 91, 92). Because hyperglycemia, dyslipidemia, low grade inflammation, not enough insulin secretion, reduced Ca2+ and K+ channel activities, and cytoskeleton changes are present in type 2 DM patients and animal models (10, 11, 44, 68), we examined the cardiac cell stiffness by using FN-coated AFM probes. The cell stiffness or elasticity was 19% increase in db/db cardiomyocytes compare to cardiomyocytes from non-db mice (in 37.9 ± 1.1 kPa in db/db mice vs 32.5 ± 1.2 kPa in non-db mice. Figure 8). These results indicated FN enhanced adhesion force and cell membrane elasticity in CVM from type 2 DM.

## Discussion

The main goal of present study was to determine the characteristics whereby extracellular matrix protein FN modulates cardiac muscle contractility in type 2 diabetes mellitus (DM) mice (db/db mice). Cardiac muscle contractility and [Ca2+]i were significantly impaired in Type-2 DM. ECM protein FN markedly increased cardiac muscle contractility and [Ca2+]i in type 2 DM. The increased effects caused by FN were greater in ventricular muscles from db/db mice than from non-db mice. FN might partially compensate for the impaired cardiac function during development of type 2 DM. Cell adhesion force and cell membrane stiffness were increased in cardiomyocytes from db/db mice in the presence of FN. These results demonstrate that FN acts increase ventricular myocyte force production and that the underlying mechanisms involve an increase in [Ca^2+^]_i_, and changes in myofilament activation processes such as Ca^2+^ sensitivity, structure cytoskeleton protein and crossbridge activation in the myocardium.

Cardiac contractile dysfunction characterized by both systolic and diastolic dysfunction in type 2 DM. Many factors may contribute to the development and progression of cardiac dysfunction in diabetes mellitus, including increased heart working efforts with decreased heart/body weight ratio, increased interstitial fibrosis, decreased intracellular Ca2+ handling, defective glucose transport, structural alterations in the form of microangiopathy, altered contractile filament properties, and/or lipotoxicity affecting both passive and active contractile properties of the heart (5, 6, 8–11, 13, 26, 36, 47, 57, 73, 83, 88). Systolic and diastolic myocardial dysfunction has been reported in a highly selected group of uncomplicated type 2 diabetic patients by using echocardiography (33). In the present study, we observed a decreased contractile function with decreased [Ca2+]i, increased body weight, waist size and heart weight, decreased heart/body weight ratio and elevated fasting blood glucose in 12-20 week-old db/db mice. we have also observed the rate of force generation (+dF/dt and −dF/dt) significant decreased with prolonged relaxation and [Ca2+]i decay in isolated papillary muscle from db/db mice. In addition, these data also suggested a possible defect in SR calcium release and uptake through sarcoendoplasmic reticular Ca2+ - ATPase (SERCA). Belke et al, used an intraventricular balloon (pressure work only) in working whole heart, observed not only a significant decrease in dP/dt in diabetic mice, but also a lower extent of pressure development. In isolated cardiac myocytes, this study also observed that the calcium transients were smaller in diabetic myocytes. Our results were supported by this study (11). It has been reported that small decreased SERCA pump activity, impaired ryanodine receptor function, decrease in sympathetic nerve activity, or increase of SERCA inhibitory protein phospholamban (PLB) in diabetic animal models (11, 31, 48, 83). All of these effects would increase the inhibitory action of PLB on SERCA activity and result in [Ca2+]i decay prolonged and slower relaxation in type 2 DM. FN compromised the impaired ventricular muscle contractility probably through increase of the phosphorylation of PLB on Ser16, the ratio of p-PLB/PLB, and the activity of L-type Ca2+ channel (94).

FN causes increased adhesion force might be because of increased integrins expression, integrin glycol-structure changes or divalent cations changes in integrin binding head induced by hyperglucose. In human umbilical vein endothelial cells, it has been observed that high glucose concentrations (30 mM) induced increased expression in FN, the FN specific integrin receptor α5β1, as well as FN, laminin, and collagen specific integrin receptor α3β1. Microvessels from diabetic patients showed increased immunostaining for β1 integrin. Overexpression of integrins correlated with increased cell attachment to exogenous fibronectin and laminin as well as to complex matrix (59). CD11a (αL), CD11b (αM), and/or CD18 (β2)-integrin levels were increased on the surface of neutrophils from type 1 and type 2 DM. The increases of αL/β2 and αM/β2 correlated with the enhanced functional adhesiveness of diabetic neutrophils to rat endothelial cell monolayers (2, 7, 61). In addition, blocking α4 integrin signaling can ameliorate the metabolic consequences of high-fat diet–induced obesity (25). Anti-integrin α4 (i.e. α4β7) treatment resulted in a significant and long-lasting suppression of the diabetic disease. However, that anti-integrin β2 treatment failed to prevent onset of type 1 DM (106, 107). It has been reported that abnormal level of metals in diabetes, such as Mg, Ca, Fe, and Cu, can cause heart disease. Treatment with metal supplements or metal chelators will decrease matrix protein and β1 integrin expression, and improve heart function in diabetes (18, 52, 65). The relationship between metal ion binding affinity and integrin ligand binding affinity has been reported (89, 103). Mg2+ uniformly facilitates and Ca2+ generally inhibits integrins with their ligands (3). Thermodynamic binding properties between FN and α5β1 integrin vary in relation to locally applied loads and divalent cations concentrations (Mg2+ and Ca^2+^) by using AFM (82). Despite extensive studies of the role of metal ions in integrin function, the possibility that metal ion binding to integrins regulated by the activation state of the integrins has not been discovered. One reason is probably that integrins contain multiple metal-binding sites, making such a determination in the native integrin difficult (104).

It has been reported that imbalance in the production and degradation of ECM proteins (e.g. FN and collagen) may lead to structural alteration including basement thickening and ECM protein deposition in the tissues as chronic DM complications (70, 93). FN expression was increased by type-1 DM and high concentration of glucose. The latter has a dose-dependent upregulation effects on FN expression. Inhibition of protein kinase C, protein kinase B, or mitogen-activated protein kinase leads to downregulation of FN expression in high glucose group (16, 47).

The cell stiffness was 19% increase in cardiomyocytes from db/db mice. Diabetes does not only impair contractile proteins but also impair structure cytoskeleton proteins. Biopsies in diabetic heart demonstrated an increase in contractile protein glycosylation (77). It has been reported that increased expression of cytoskeletal proteins, α-smooth muscle actin and vimentin, and extracellular matrix proteins within the kidneys of diabetic rats (63). In Type 2 DM, neutrophil surface αMβ2 integrin (CD11b/CD18) expression is increased and is associated with impaired actin polymerization and increased cytoskeletal phosphotyrosine (2). In Type 2 diabetes, neutrophil actin polymerization in response to increasing tyrosine phosphorylation is impaired despite an overall increase in cytoskeletal phosphotyrosine. Impaired neutrophil cytoskeletal function is also associated with decreased calcium signaling in Type 2 DM. Correction of the cytoskeletal defect in Type 2 diabetes may have a positive effect on cardiovascular morbidity and mortality. (1). In addition, ECM proteins (e.g. CN and FN) deposits compromise tissue compliance and causing myocardial stiffness (8, 14). Jin et al reported that increased stiffness and adhesion force between AFM tip and cell memberane of erythrocytes from type 2 DM patients (43).

In conclusion, our data indicate that FN increased contractility and Ca2+ concentration, and increased mechanical properties on impaired diabetic heart. Further studies will be required to clarify whether FN effect is only temporary compensation, and which integrin involved in the mechanisms by which these adhesive changes occur and to determine if such changes in adhesion can be modulated on a beat-to-beat basis in cardiomyocytes (12). To our knowledge, this is the first report to advance and provide support for the hypothesis that integrin adhesive properties are modulated in diabetic cardiomyocytes by using nanotechnology of atomic force microscopy. Heart disease is the leading cause of death (>65%) among diabetic patients in United States. Diabetic cardiomyopathy is a disease that occurs in the absence of vascular disease, e.g. atherosclerosis. About 50% diabetic patients with or without complications (e.g. cardiovascular diseases) cannot be controlled by using current marketed various medications (51). Management of diabetic cardiomyopathy and cardiac hypertrophic patients should not only simply focus on normalization of blood glucose and pressure, but also have to target the adverse structural and functional feature that cause ventricular remodeling including CVM and non-CVM compartments by diabetes (20, 32, 42). The integrin ligands might be the potential drug candidates for the treatment and/or prevention of human diseases including diabetes and cardiovascular diseases (7, 19, 22, 37, 87, 96, 99). Thus, understanding the mechanisms that integrin modulated biomechanical forces to the activation of stress pathways will give insight into the biology of heart diseases.

## ACKNOWLEDGEMENTS

This work was supported by Texas A&M Health Science Center Research Development & Enhancement Award Program 244441-20702 to WX and National Institutes of Health grants R21 EB003888-01A1 to MM.

